# Use of Continuous Traits Can Improve Morphological Phylogenetics

**DOI:** 10.1101/121343

**Authors:** Caroline Parins-Fukuchi

## Abstract

The recent surge in enthusiasm for simultaneously inferring relationships from extinct and extant species has reinvigorated interest in statistical approaches for modelling morphological evolution. Current statistical methods use the Mk model to describe substitutions between discrete character states. Although representing a significant step forward, the Mk model presents challenges in biological interpretation, and its adequacy in modelling morphological evolution has not been well explored. Another major hurdle in morphological phylogenetics concerns the process of character coding of discrete characters. The often subjective nature of discrete character coding can generate discordant results that are rooted in individual researchers’ subjective interpretations. Employing continuous measurements to infer phylogenies may alleviate some of these issues. Although not widely used in the inference of topology, models describing the evolution of continuous characters have been well examined, and their statistical behaviour is well understood. Also, continuous measurements avoid the substantial ambiguity often associated with the assignment of discrete characters to states. I present a set of simulations to determine whether use of continuous characters is a feasible alternative or supplement to discrete characters for inferring phylogeny. I compare relative reconstruction accuracy by inferring phylogenies from simulated continuous and discrete characters. These tests demonstrate significant promise for continuous traits by demonstrating their higher overall accuracy as compared to reconstruction from discrete characters under Mk when simulated under unbounded Brownian motion, and equal performance when simulated under an Ornstein-Uhlenbeck model. Continuous characters also perform reasonably well in the presence of covariance between sites. I argue that inferring phylogenies directly from continuous traits may be benefit efforts to maximise phylogenetic information in morphological datasets by preserving larger variation in state space compared to many discretisation schemes. I also suggest that the use of continuous trait models in phylogenetic reconstruction may alleviate potential concerns of discrete character model adequacy, while identifying areas that require further study in this area. This study provides an initial controlled demonstration of the efficacy of continuous characters in phylogenetic inference.

---

The development and widespread adoption of statistical phylogenetic methods has revolutionized disparate disciplines in evolutionary biology, epidemiology, and systematics. Studies utilizing maximum-likelihood (ML) and Bayesian approaches have become the preferred means to analyse molecular data, largely eclipsing parsimony and distance methods. Despite this, approaches which draw inference from morphological data have remained comparatively underdeveloped (but see relevant discussion and citations below). As a result, non-probabilistic tree inference methods have continued to be employed for the phylogenetic analysis of morphological characters. Nonetheless, several landmark advances in the development of statistical morphological phylogenetic methods have demonstrated the benefits of further developing this framework. This will be particularly important in the near future as burgeoning approaches enabling the rapid collection of morphological data may begin to outstrip methods through which to analyse them (Chang and Alfaro 2015b,a). This may significantly alter and enhance our view of the tree of life, especially considering that the majority of macro-organisms, represented by fossil taxa, can only be analysed from their morphology.

A foundational contribution in morphological phylogenetics has been the Mk model of discrete trait evolution (Lewis 2001). This is a version of the Jukes-Cantor model of nucleotide substitution generalised to accommodate varying numbers of character states (Jukes and Cantor 1969). Extensions to this model accommodate for biased sampling of parsimony informative characters (Lewis 2001), rate heterogeneity between sites (Wagner 2012), and asymmetric transition rates (Ronquist and Huelsenbeck 2003; Wright *et al.* 2015). The deployment of this model has demonstrated the utility of statistical approaches to morphological phylogenetics. Such approaches improve estimates of uncertainty over non-probabilistic approaches, enable a clearer statement of modelling assumptions, and enable branch length estimation. This has enabled a better understanding of much of the fossil tree of life (Dávalos *et al.* 2014; Pattinson *et al.* 2014; Dembo *et al.* 2015). These approaches have also enabled the application of tip dating methods to the combined analysis of extinct taxa represented by morphological data with extant taxa (Nylander *et al.* 2004; Ronquist *et al.* 2012). These total evidence tip dating methods have been widely used since their introduction, and are implemented in the BEAST (Bouckaert *et al.* 2014) and MrBayes (Ronquist and Huelsenbeck 2003) packages. These have more clearly resolved the timing of species divergences and relationships between fossil and living taxa (Wiens *et al.* 2010; Wood *et al.* 2012; Lee *et al.* 2013, 2014, but see Arcila *et al.* (2015)). Overall, probabilistic approaches to morphological phylogenetics appear to represent an improvement in accuracy compared to cladistic methods, and are indispensable in their distinct ability to allow the estimation of branch lengths and evolutionary rate. The benefits of a statistical total-evidence framework as applied to fossil taxa will only become clearer as more data become available and improved methods are developed (Pennell and Harmon 2013; Lee and Palci 2015).

Despite the these strides, discrete character models represent an imperfect solution in their current usage. Although Bayesian inference under Mk appears to outperform parsimony under certain conditions, error increases at high evolutionary rates (Wright and Hillis 2014). Also, under many circumstances, phylogenetic inference under the Mk model includes imprecision and uncertainty, both in simulations (O’Reilly *et al.* 2016; Puttick *et al.* 2017) and empirical studies (Lee and Worthy 2012; Dembo *et al.* 2015). Previous researchers have also expressed concerns over the efficacy of model-based approaches in the presence of missing data (Livezey and Zusi 2007; O’leary *et al.* 2013). However, these have been assuaged and any issues arising from missing data are likely not specific to probabilistic approaches (Wright and Hillis 2014; Guillerme and Cooper 2016). Another potential issue is the lack of clarity in interpreting the Mk model biologically. Although transition rates have a strong theoretical and empirical basis in population genetics, their significance beyond serving as nuisance parameters is less straightforward when applied to morphological data. Discrete morphological characters may not undergo change in a manner analogous to nucleotides, which are well understood to alternate between states repeatedly. Conversely, many characters used for phylogenetic inference consist of single, parsimony informative directional changes between taxa (Klopfstein *et al.* 2015). It is unclear how adequately discrete Markov models describe such variation. The Mk model itself does not accommodate directional evolution, and previous researchers have questioned the adequacy of existing discrete character models (Ronquist *et al.* 2016). This is particularly important when considering the importance of branch lengths in total evidence tip dating methods discussed above, but may also be expected to mislead inference of topology.

Aside from the modelling concerns discussed above, discrete morphological characters present a non-trivial set of challenges to phylogenetics that are distinct from those possessed by molecular data. Perhaps foremost among these is disagreement between researchers in the categorisation, ordering, and weighing of discrete character states (Farris 1990; Hauser and Presch 1991; Pleijel 1995; Wilkinson 1995). Despite extensive discussion among comparative biologists, the interpretive nature of the process of character coding has continued to leave major palaenotological questions unresolved (Upchurch 1995; Wilson and Sereno 1998; Bloch and Boyer 2002; Kirk *et al.* 2003).

Use of continuous characters may help to address some of the concerns with discrete traits discussed above. They can be collected more objectively than qualitative observations and do not require ordering of states. Their use in phylogenetic inference has been discussed among the earliest advancements in statistical phylogenetics (Cavalli-Sforza and Edwards 1967; Felsenstein 1973), and their phylogenetic informativeness has been demonstrated empirically (Goloboff *et al.* 2006; Smith and Hendricks 2013). Still, the use of continuous characters for the inference of phylogenetic topology has remained uncommon, with methods for their use in phylogenetics remaining relatively poorly examined beyond the foundational works referenced above. Although many palaeontological studies incorporate continuous measurements, they are binned into categories and analysed as discrete. However, since fossil data are often scarce, it may be beneficial to maximise the amount of information gleaned from available specimens by representing such variation in its entirety.

Another potential benefit to inferring phylogeny from continuous characters is the wealth of models developed in phylogenetic comparative methods to describe their evolution. Most comparative models of continuous trait evolution belong to the Gaussian class, which are also well utilized in disparate fields such as physics, economics, and engineering. In comparative biology, they are used to describe stochastic Markovian movement through continuous trait space along continuous time. This class of models includes Brownian motion (BM) (Felsenstein 1973, 1985; Gingerich 1993), Ornstein-Uhlenbeck (OU) (Hansen 1997; Butler and King 2004; Beaulieu *et al.* 2012), and Lévy processes (Landis *et al.* 2013). Under BM, evolution is described as a random walk, with phenotypic change being normally distributed with a mean displacement of zero, and variance *σ*^2^. OU models expand upon this by introducing terms producing a stabilizing force which stabilizes movement around an optimal trait value, while Lévy processes contain terms producing saltational jumps in character space, interspersed either by BM diffusion or stasis. Two major benefits to Gaussian models in phylogenetics are their relatively straightforward interpretability and the relative ease of deriving mathematical extensions to describe a range of biological processes.

Given the existence of well understood and clearly interpretable models describing their evolution, the use of continuous traits may offer several advantages over discrete characters in phylogenetic inference. However, their behaviour is not well understood when applied to the inference of phylogenetic topology, and so further investigation is needed. In addition, there are potential hurdles to their efficacy. Possibly foremost among these is the widespread covariance between continuous measurements that is expected through both genetic and morphometric perspectives (Lynch *et al.* 1998; Uyeda *et al.* 2015; Adams and Felice 2014). Nevertheless, the expected magnitude in covariance among continuous morphological measurements and the robustness of phylogenetic methods to this violation is not known. Furthermore, it is also generally reasonable to expect evolutionary covariance between nucleotide sites, and phylogenetic methods that do not accommodate for this are routinely applied to molecular data.

In this study, I carry out simulations to compare the relative performance of binary discrete and continuous characters at reconstructing phylogenetic relationships. Simulations of continuous characters were designed to reflect a range of scenarios that may influence accuracy including overall evolutionary rate and matrix sizes. I also conduct inference on continuous traits that have undergone correlated evolution, an important violation to single-rate BM thought to be widespread in continuous character evolution.

## METHODS

### Simulations

I generated a set of 100 pure birth trees using the Phytools package (Revell 2012) package in R (R Core Team 2016), each containing ten taxa. All trees were ultrametric and generated with a total length of 1.0 units for consistency in parameter scaling for trait simulations (Fig. 1). These trees were used to simulate continuous characters evolving along an unbounded BM process, again using Phytools. This is a Markovian process in continuous time where the variance of the process can increase infinitely through time. This differs from the BM *σ*^2^ parameter, which gives the variance in the amount of character displacement at each draw, effectively describing the magnitude of the random BM walk or a rate of character displacement. To assess performance across several biological scenarios, traits were simulated at *σ*^2^ parameterizations of 0.05, 0.5, 1.0, 1.5, and 3. Since the process under which traits were simulated is unbounded, phylogenetic signal is expected to remain consistent across rates (Revell *et al.* 2008), but different rates were chosen to illustrate this consistency and to provide even comparison to discrete trait simulations. Discrete characters were simulated in the Phytools package (Revell 2012) under an Mk model with homogeneous transition probabilities. Traits were generated at transition rates 0.05, 0.5, 1.0, 1.5, and 3. All character matrices were generated without rate heterogeneity, and include invariable sites (ie. no acquisition bias).

**Figure 1:**
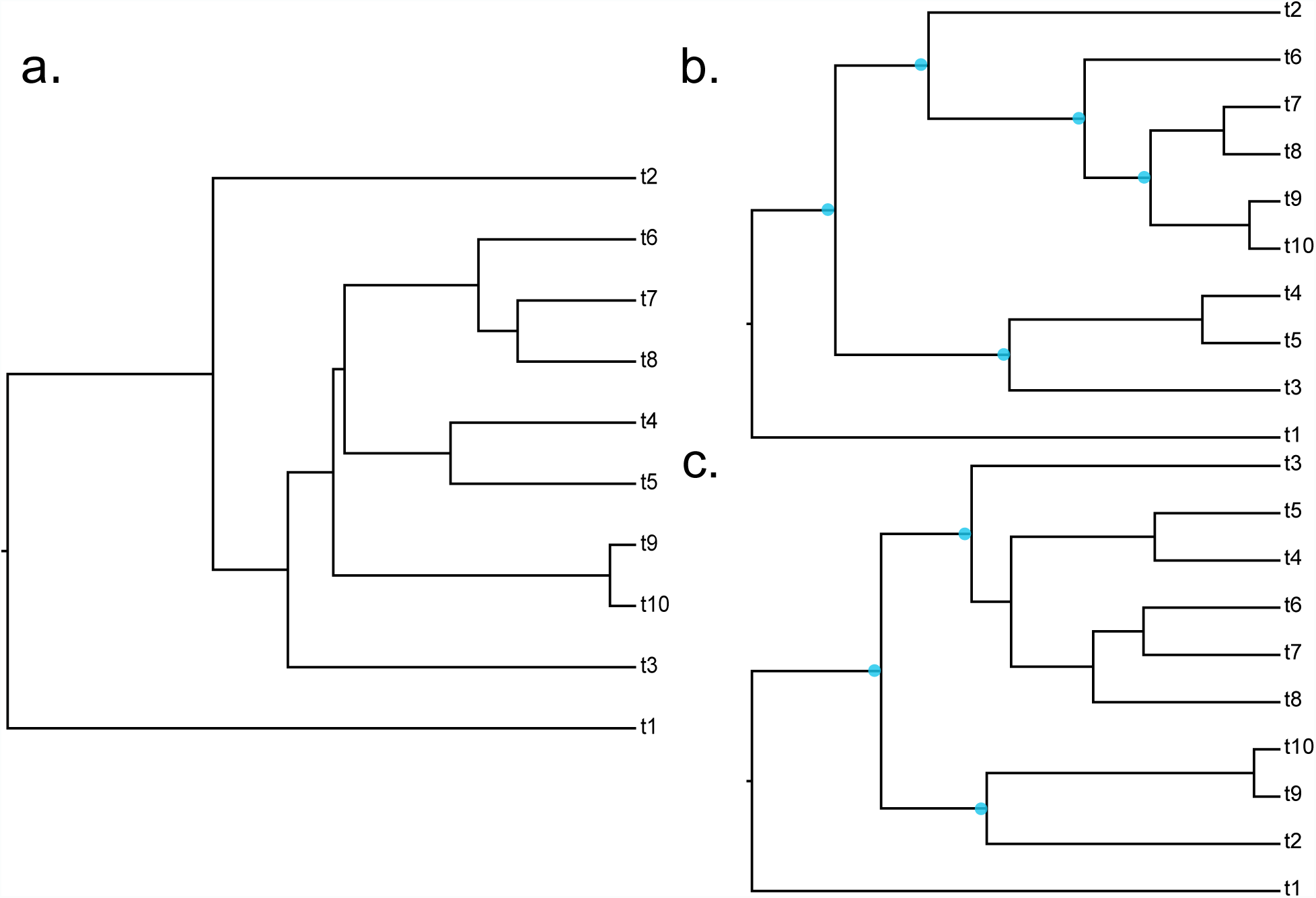
**a.** Exemplar true simulated tree. **b.** Tree inferred from 20 discrete characters simulated under Mk from true tree. **c.** Tree inferred from 20 continuous characters simulated under Brownian motion. Blue dots denote incorrect bipartitions.

Matrices containing 500 traits were generated and randomly subsampled to create smaller sets of 20 and 100 characters to reflect a range of sampling depths. These were chosen because many published morphological matrices fall within this range. The subsampled matrix sizes were chosen to represent reasonably sized palaeontological datasets, while the 500 trait matrices were tested to assess performance when data are abundant. While such large datasets are uncommon in morphology, several studies have produced character matrices of this size, and for continuous characters, it may be feasible to generate such large datasets from morphometric data.

I also simulated continuous characters under an OU model parameterised without directional drift (*θ* = 0), and with the stabilizing (*α*) parameter set to yield the same phylogenetic half-life present in the binary Mk model used for comparison. For OU continuous characters, phylogenetic half-life is defined by:

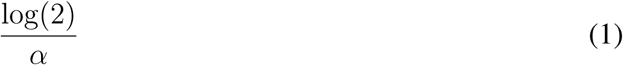

and for binary discrete characters as:

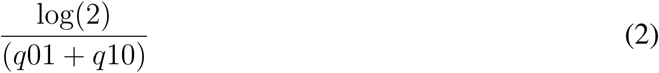

With *q*01 and *q*10 corresponding to the respective transition rates between binary character states.

When phylogenetic half-life is set to be equal, phylogenetic constraint should be the same between both sets of characters in the sense that they reach saturation over the same timescale. This comparison examines whether either data source performs inherently better when phylogenetic signal is held constant. These data were generated in matrices of 100 traits at an evolutionary rate of 0.5. Because the phylogenetic information content of both sets of constrained traits should be the same, both sets are expected to perform similarly. Nevertheless, this comparison provides a control by assessing whether unknown differences in the behaviour of each model (or other properties of each method) themselves lead to any differences in reconstruction accuracy.

Data were also generated under a correlated BM process to mimic inference in the presence of multidimensionality. These datasets were constructed at covariance strengths of 0.1, 0.5, and 0.9 and covarying dimensions of 5 and 25 traits. These were chosen to represent situations where traits range from being loosely to tightly correlated to each another, and where the number of correlated dimensions is large to small. Although differing, these values were chosen to loosely follow the scheme of Adams and Felice (2014).

### Estimation of Phylogenies and Reconstruction Accuracy

I estimated Bayesian phylogenetic trees under a single rate BM model for all sets of continuous characters using RevBayes (Höhna *et al.* 2016). Trait likelihoods were computed after Felsenstein (1973, 1985). MCMC simulations were run for 150,000-1,000,000 generations and checked manually for convergence using Tracer v1.6 (http://tree.bio.ed.ac.uk/software/tracer/). Runs were accepted when the effective sample size (ESS) for logged parameters exceeded 200. Trees were inferred from discrete data in MrBayes version 3.2.6 (Ronquist and Huelsenbeck 2003), simulating for 1,000,000 generations. Different programs were used because, while MrBayes remains the standard in the field for Bayesian phylogenetic inference, its current version does not implement likelihood functions for continuous character models. So the continuous character approach needed to be developed in RevBayes, however, I preferred to remain with the standard and proven implementation where possible. For both continuous and discrete characters, I incorporated a birth-death prior on node heights. This was done to enable an even comparison of branch lengths obtained through both methods that are scaled to time. Example configuration files for RevBayes and MrBayes analyses are provided as supplementary data. Tree distributions were summarized using TreeAnnotator version 2.4.2 (Rambaut and Drummond 2013) to yield maximum clade credibility (MCC) topologies. MCC trees maximize the posterior probability of each individual clade, summarizing across all trees sampled during MCMC simulation. Once summarised, all trees were rescaled to match inferred tree lengths to the true trees using Phyx (https://github.com/FePhyFoFum/phyx).

I assessed topological accuracy from simulated trait data using the symmetric (Robinson-Foulds) distance measure (Robinson and Foulds 1981), giving the topological distance between true trees and inferred trees. Symmetric distance is calculated as a count of the number of shared and unshared partitions between compared trees. As such, the maximum symmetric distance between two unrooted trees can be calculated as *2(N-3)*. These values were then scaled to the total possible symmetric distance for interpretability. Additionally, I measured error in branch length reconstruction using the branch length distance (BLD) (Kuhner and Felsenstein 1994). This is calculated as the sum of the vector representing the individual differences between the branch lengths of all shared bipartitions. The scale of this value depends on the lengths of the trees under comparison. If trees of different lengths are compared, BLD can be very high. However, in this study, all trees are scaled to a root height of 1 to allow comparison of topological and internal branch length reconstruction error. All distances were calculated using the DendroPy Python package (Sukumaran and Holder 2010). Summary barplots were constructed using ggplot2 (Wickham 2016).

## RESULTS

### Unconstrained and Independently Evolving Continuous Traits

Topological reconstruction error is lower overall for trees estimated from continuous characters than from binary discrete (Fig. 2a, Supp. Fig, 1a). For discrete characters, symmetric distance increases significantly at high evolutionary rates, likely due to saturation and loss of phylogenetic signal. Distance also increases in discrete characters when rate is very slow, due to lack of time for phylogenetic signal to develop. This pattern is similar to that recovered by (Wright and Hillis 2014) in their test of Bayesian inference of Mk, which revealed highest topological error at very low and high rates. As expected, continuous characters perform consistently across rates because saturation cannot occur, even at very fast rates. Because of the differing sensitivities of each data type to evolutionary rate, topological error should also be compared using the most favourable rate class for discrete characters, 0.5 substitutions per million years (Fig. 2b, Supp. Fig. 1b). Even at this rate, continuous reconstruction performs more consistently than discrete, with error more tightly distributed around a slightly lower mean. A likely explanation is that discrete characters retain less information that continuous characters. The small state space of the binary character model likely causes phylogenetic signal to become saturated more quickly at fast rates, and develop too slowly at slow rates than multi-state characters. BM and Mk appear to perform fairly similarly in reconstructing branch lengths (Fig. 2; Supp. Fig. 1). The pattern across rates and matrix sizes are very similar between BLD and symmetric distances, with the fastest rates producing the most error. This likely results from increased saturation at fast rates, causing underestimation of hidden character changes.

**Figure 2:**
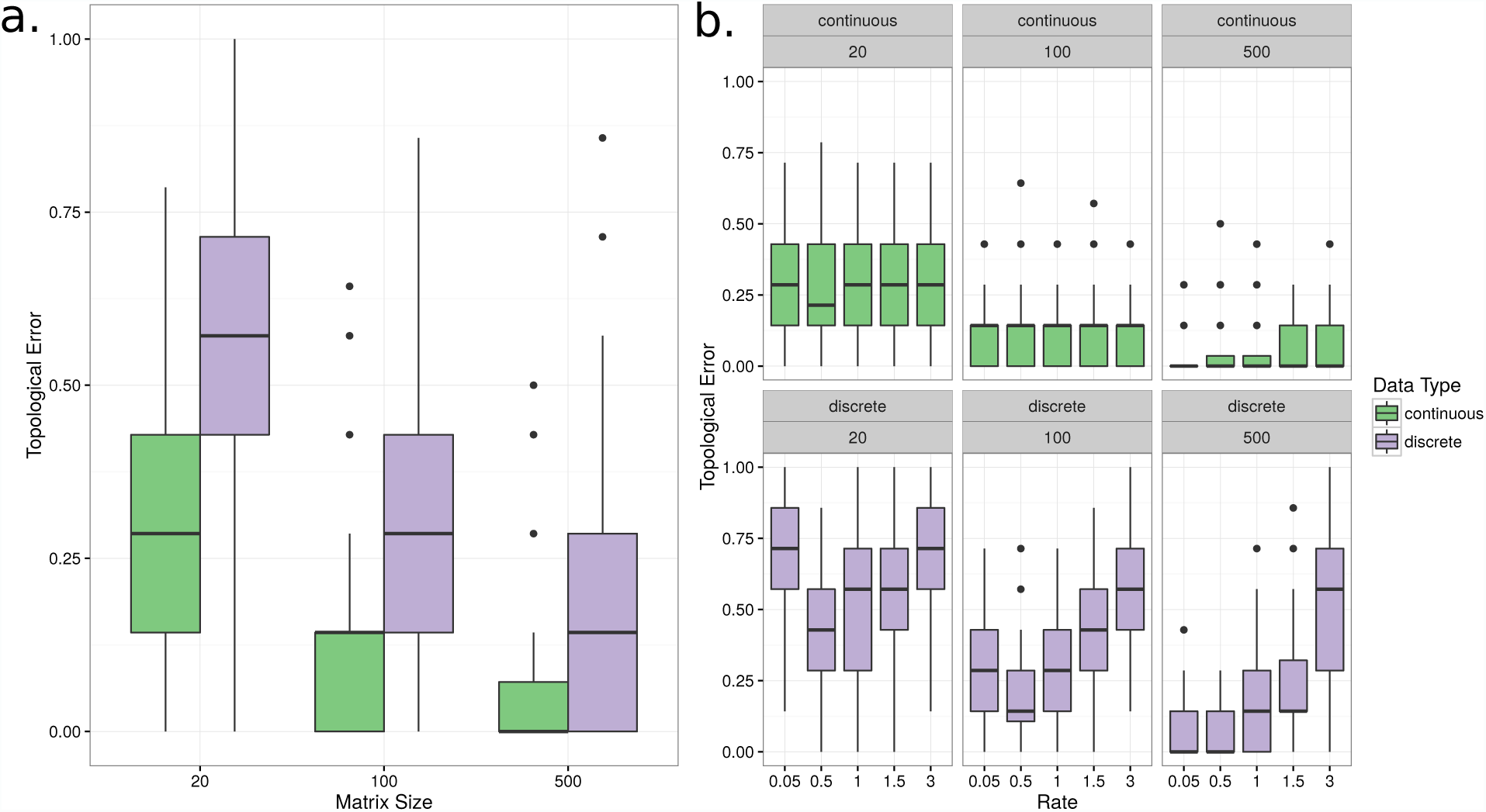
Topological error calculated as the proportion of maximum symmetric distance across trees estimated from independently evolving continuous characters. **a.** Error averaged across all rates except for the highest rate category, which resulted in the highest error when inferring under Mk. **b.** Error across all matrix sizes and rates.

Matrix size has a major impact on tree reconstruction accuracy. Estimations from both discrete and continuous traits improve substantially at each increasing matrix size (Fig. 2). Estimates from 20-character matrices possess fairly high error in both data types, with approximately 1 in 5 bipartitions being incorrectly estimated from continuous characters, and 2 in 5 incorrectly being incorrectly estimated from discrete data. Increasing matrix size to 100 traits improves accuracy significantly, with both data types estimating approximately 1 in 10 bipartitions incorrectly. Although at several rates, mean symmetric distance compared between data types is close, continuous characters tend to be less widely distributed, and thus appear to reconstruct trees with more consistent accuracy. When matrix size is increased to 500 characters, both continuous and discrete characters are able to recover phylogeny with very high accuracy, except for at very fast rates, where discrete characters estimate approximately half of all bipartitions incorrectly on average.

### Continuous Traits Evolving Under Selective Constraint

Phylogenies inferred from continuous traits simulated under an OU model achieve virtually identical performance to binary discrete characters simulated under the same phylogenetic constraint (Fig. 3). Both sets of characters display a very similar range of error, with approximately 15% of bipartitions estimated incorrectly on average. This result demonstrates that any performance increases observed for continuous traits over discrete traits result from differences in realised phylogenetic information.

**Figure 3:**
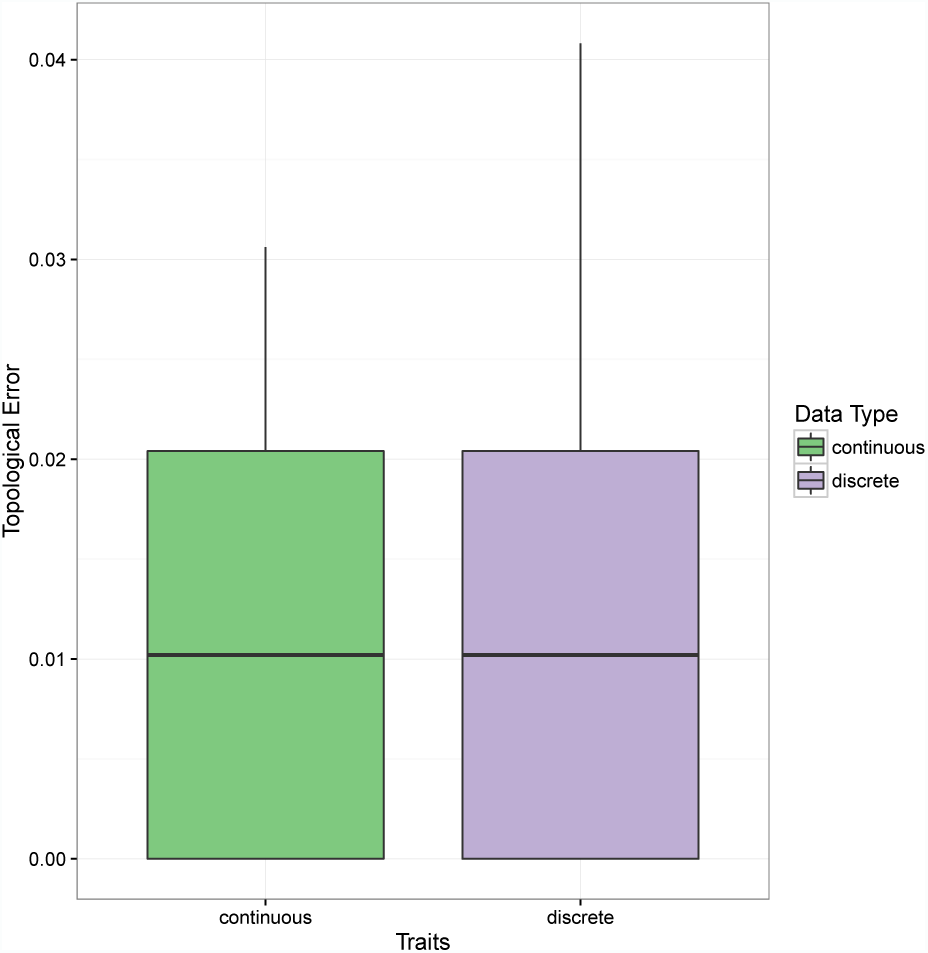
Topological error achieved after reconstructing trees from discrete traits simulated under Mk at rate 0.5, and single rate Ornstein Uhlenbeck at rate 0.5 with no directional drift and constraint set equal to the discrete characters.

### Covarying Continuous Characters

Tree inference under BM appears relatively robust to the violation of co-evolving continuous characters. Although error is recognisably greater with strong covariance and many trait dimensions, symmetric distance is remains close to values from uncorrelated traits at lower covariance strengths and/or fewer trait dimensions (Fig. 4). When correlated traits are of low dimensionality and covariance strength, reconstruction appears to be nearly as accurate as uncorrelated traits, with all bipartions estimated correctly on average. As covariance strength and dimensionality are increased to intermediate values, topological error increases such that between 0 and 17% of bipartitions are estimated incorrectly, with a wider distribution than is present at the lowest values. Accuracy is most diminished when covariance is strongest and dimensionality is largest, with most reconstructions estimating between 17-29% of bipartitions incorrectly. Although statistical significance cannot be estimated for BLD and symmetric distance, estimation under low to intermediate trait covariance appears at least qualitatively similar, albeit slightly worse, to uncorrelated continuous and binary discrete characters. The decreases in accuracy observed can likely be attributed to the decrease in total information content caused by covariance. This reduces the effective amount of data from which to draw inference. This is reflected in the results, with higher covariances and dimensionalities reconstructing trees with a similar magnitude of error as is shown for the 100 character datasets.

**Figure 4:**
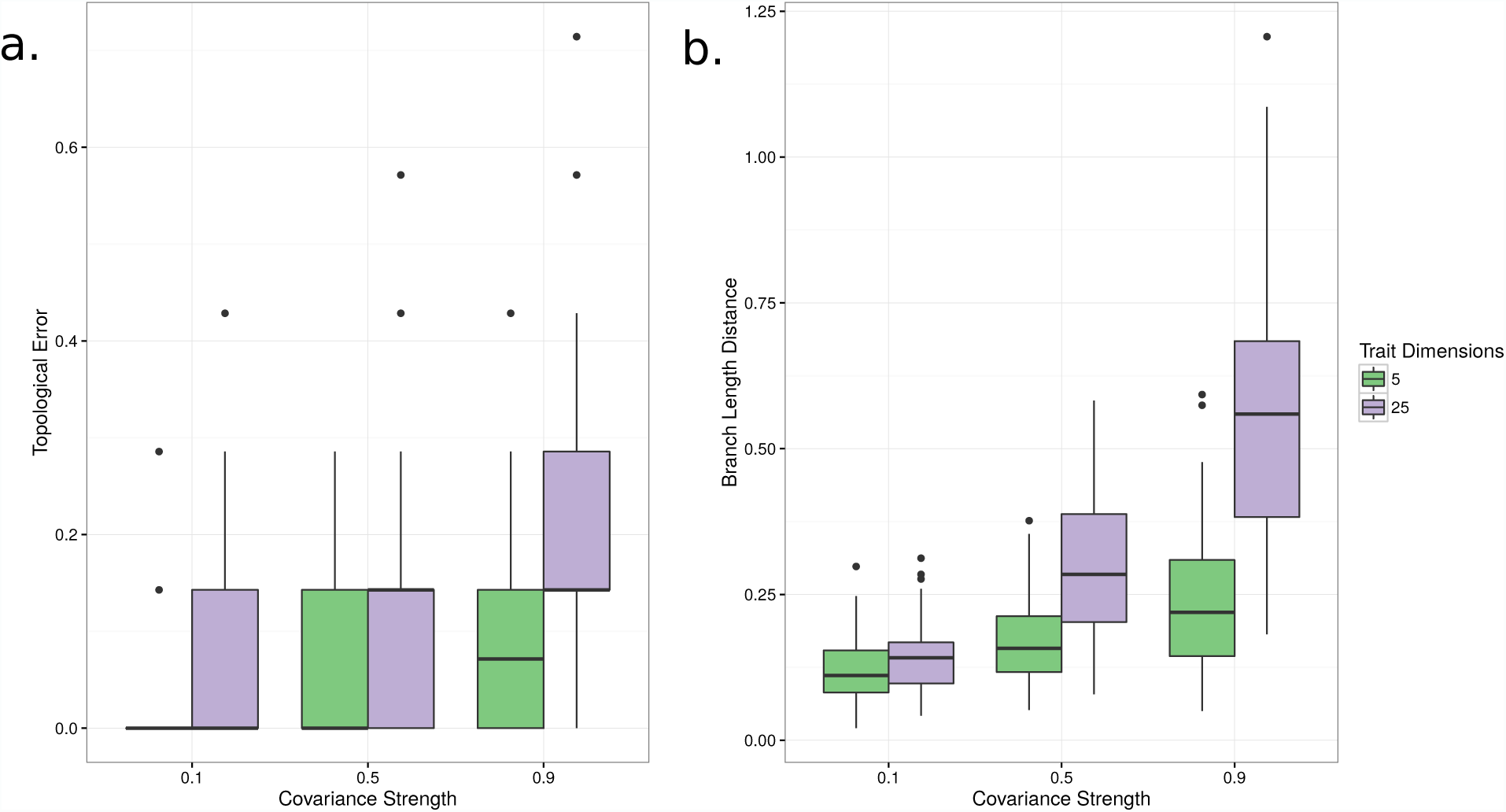
**a.** Topological error, calculated as proportion of maximum symmetric distance across trees estimated from covarying continuous characters. **b.** Branch length distance (BLD) across trees estimated from covarying continuous characters. Dimensions refers to the number of traits within covarying blocks. Covariance strength refers to the strength of the correlation between covarying characters, with a value of 0 describing to complete independence and 1 describing perfect correlation.

**Figure 5:**
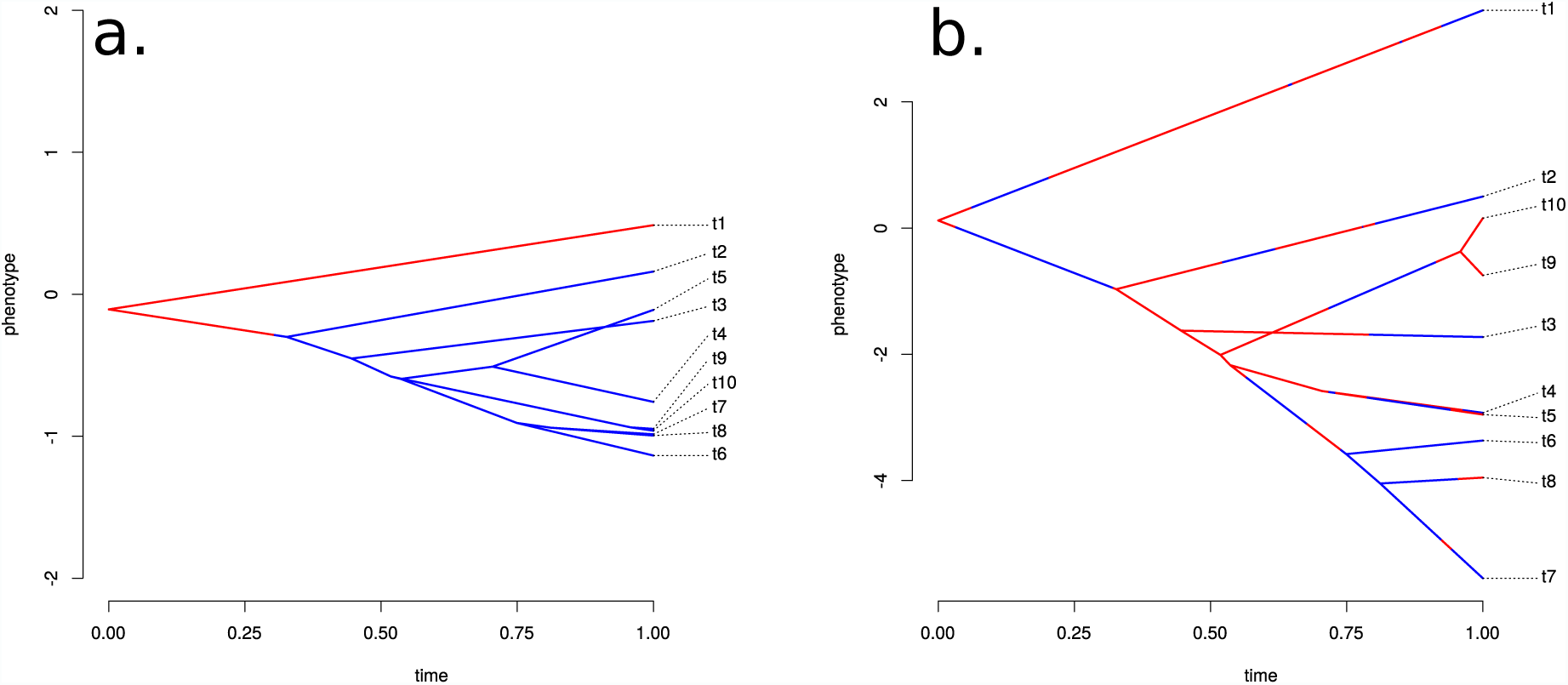
Discrete and continuous characters simulated **a.** at slow evolutionary rate and **b.** fast evolutionary rate. Y axis represents continuous phenotype. Changes in colour represent changes in discrete character state. Note how continuous characters retain phylogenetic signal at fast rates, while discrete characters saturate. Figure drawn using phytools (Revell 2012).

## DISCUSSION

The results demonstrate that phylogenetic reconstruction from continuous trait data can provide a reasonable supplement or alternative to inference from discrete characters. Continuous characters that are unconstrained and unbounded in their evolution outperform discrete characters, and perform equally well when constrained by selection. The unconstrained traits’ resilience to high evolutionary rate is expected, because continuous characters evolving under an unbounded and unconstrained BM process will continue to increase in variance through time. Therefore, such characters are able to retain phylogenetic information at high evolutionary rates that may cause rampant saturation in discrete characters (Fig. 4). Further work is needed in this area to investigate the extent to which continuous characters are bounded and constrained in their evolution relative to discrete characters. This will be especially important moving forward, as temporal variation in evolutionary regimes and model parameters can interact in complex ways, sometimes extending the maintenance of phylogenetic signal through time (Revell *et al.* 2008). Although continuous characters in empirical are undoubtedly constrained in their evolution, the added information contained in continuous character datasets may lessen the extent of saturation relative to discrete characters in practice.

The demonstration that performance becomes equal when the amount of phylogenetic constraint is held constant between both data sources identifies the major source of the performance increase observed in unconstrained BM traits compared to discrete traits. The average amount of phylogenetic constraint exhibited by discrete and continuous traits, however, is not well understood in empirical datasets. Conversely, the susceptibility of discrete traits to the loss of phylogenetic signal at high evolutionary rates and deep timescales has long been recognised (Hillis and Huelsenbeck 1992; Yang 1998). Although this effect is understood to affect molecular data, discrete morphological datasets may possess increased susceptibility to this effect because of the frequent use of binary character coding schemes. Discrete characters constrained to fewer states increases signal loss at high evolutionary rates due to increased levels of homoplasy, saturation, and lower information content overall (Donoghue and Ree 2000). The extent to which continuous traits are constrained in their evolution on average is not well understood. However, the results here suggest that researchers would benefit in treating continuous traits as such and inferring phylogenies under continuous trait models in order to maximise usable information contained in datasets.

My results demonstrate that the fundamental issues in comparing continuous and discrete traits are state space, selective constraint, and evolutionary boundedness. When selective constraint in continuous characters occurs at levels which restrict phylogenetic signal with the same strength as binary characters, reconstruction accuracy is predictably equal. Nevertheless, it is unclear the extent to which phylogenetic half-life in continuous and discrete traits tends to differ in empirical datasets. Continuous characters may be expected to commonly evolve under some manifestation of selective constraint, but it is unclear whether such effects typically mask phylogenetic signal to the same extent as rapidly saturating binary traits.

Discrete traits with more than two states possess a significantly longer phylogenetic half-life than binary characters, but could be supplanted by continuous characters in many cases. Although empirical morphological datasets typically incorporate discrete characters with more than two states, these are typically fewer in number than binary coded characters. Multi-state characters are also typically discretized codings of continuous measurements. Such "discrete" traits would be susceptible to the same selective forces as their continuous counterparts, and so treatment of the multi-state partitions of morphological matrices as continuous can only increase the amount of phylogenetic information contained within datasets. The tendency of morphological matrices to be predominantly composed of binary characters should encourage further consideration of continuous traits in future empirical and theoretical studies.

Error in branch length estimation was fairly high with the 20-trait matrices but decreased substantially when matrix size was increased to 100 traits. Although BM and Mk achieve similar accuracy in estimating branch lengths in this study, careful thought should continue to be applied when relying upon Mk branch length estimates in the future. Branch length error may be higher when inferring under Mk from empirical datasets, since many discrete morphological matrices are constructed to include only parsimony informative characters. In these cases, characters are expected to have undergone only single synapomorphic changes. Although the lack of invariable sites in datasets tailored to parsimony is addressed through the ascertainment bias correction developed by (Lewis 2001), it is unclear how meaningfully the directional single character changes often observed in these datasets can inform evolutionary rates. This mode of change, which may characterise much of discrete character evolution, differs from the population dynamics of nucleotide substitution.

Although continuous traits may often follow covarying evolutionary trajectories in nature, this appears to have a relatively minor impact on reconstruction. Accuracy was only greatly lowered in the simultaneous presence of very high dimensionality and covariance strength. Offering further support to the ability of continuous characters to reconstruct phylogeny despite evolutionary covariance, Adams and Felice (2014) also report the presence of phylogenetic information in multidimensional characters, even when the number of dimensions is greater than the number of taxa. Despite these generally positive findings, it should be noted that inference may be misled if sampling is significantly biased to include relatively small numbers of strongly correlated measurements. In these cases, it would be beneficial to examine the correlation structure and information content of the dataset to assess the amount of biased redundancy in signal.

### Can Using Continuous Characters Benefit Morphological Phylogenetics?

Use of continuous traits has the benefit of reducing subjectivity in the construction of data matrices in many cases. Categorizing qualitative characters often requires subjective interpretation. However, quantitative measurements can be taken without this source of human error. This increased objectivity in the measurement of quantitative characters would expand biologists’ capacity to assess statistical uncertainty. Although the likelihood approaches to morphological phylogenetics enabled by the Mk model represent a major step in this direction, discordance in tree estimates can still be attributed to differences in qualitative categorization of variation by researchers. Translation of morphological observations into data that can be analysed can present serious complications in discrete characters. Steps such as the determination of whether or not to order states, the total number of states chosen to describe characters, and the assignment of character states can vary greatly and often yield widely different results (Hauser and Presch 1991; Pleijel 1995; Wilkinson 1995; Hawkins *et al.* 1997; Scotland and Pennington 2000; Scotland *et al.* 2003; Brazeau 2011; Simões *et al.* 2017). Continuous measurements avoid many of these issues because they can be measured, by definition, objectively and quantitatively. In addition, they may better describe variation than discrete characters. Several workers have suggested that the majority of biological variation is fundamentally continuous (Thiele 1993; Rae 1998; Wiens 2001). Although continuous characters have long been employed in phylogenetic analysis, they are generally artificially discretised, either by applying thresholds to interspecific measurements or through gross categorisations such as “large” and “small”. The major disadvantage to this approach is the loss of valuable biological information. Several researchers have condemned the use of continuous characters in phylogenetics, arguing that intraspecific variation may be too great for clear phylogenetic signal to exist (Pimentcl and Riggins 1987; Chappill 1989). However, these arguments have been largely undermined by studies demonstrating the phylogenetic informativeness of continuous measurements (Goloboff *et al.* 2006; Smith and Hendricks 2013).

The expectation of correlated evolution between continuous characters has been a major argument against their use in phylogenetic reconstruction in the past (Felsenstein 1985). However, evolutionary covariance between sites is not a phenomenon that is restricted to continuous morphological characters. Population genetic theory predicts tight covariance between nucleotide sites under many conditions (e.g. Hill and Robertson 1968; Reich *et al.* 2001; Palaisa *et al.* 2004; Schlenke and Begun 2004; McVean 2007). Such covariance has also been demonstrated among discrete characters (Pagel 1994), and so this concern is not unique to continuous measurements but is shared by all phylogenetic approaches. While it is difficult to assess the relative magnitude of sitewise covariance between continuous, discrete, and molecular data, examination of the correlation structure of traits may be more straightforward in continuous characters using standard regressional techniques. This would ease the identification of biased and positively misleading signal among continuous characters, enabling correction through common transformation approaches such as principal components analyses or by weighting likelihood calculations by the amount of overall variance contributed by covarying sets of characters.

The fundamentally continuous nature of many biological traits is supported by differential gene expression and quantitative trait loci mapping studies, which demonstrate their quantitative genetic basis (Andersson *et al.* 1994; Hunt *et al.* 1998; Frary *et al.* 2000; Valdar *et al.* 2006). Nevertheless, there remain well known instances where traits are truly discrete. Studies in evolutionary developmental biology have shown that many traits can be switched on or off in response to single genes controlling genetic cascades (e.g. Wilkinson *et al.* 1989; Burke *et al.* 1995; Cohn and Tickle 1999). Characters used in phylogenetic analysis are also frequently truly discrete, representing qualitative categories (eg., presence/absence). These traits may be incorporated as separate partitions into integrated analyses along with continuous measurements (Fig. 6). Such combined analyses can be performed in RevBayes by adding a discrete trait model, such as Mk, and discrete character data. In practice, this may improve inference from discrete characters alone, and would represent a conceptual advance in its ability to treat all available data as faithfully as is possible. Doing so may improve upon existing paradigms, which group continuous variation into multi-state discrete characters, potentially preserving more phylogenetic information. An added benefit would be the greater flexibility in modelling the evolution of such traits by making available all existing continuous trait models. An example RevBayes script for a phylogenetic analysis combining continuous and discrete characters is available in the supplement. Characters under the control of developmental expression pathways may also exhibit very deep phylogenetic signal (De Rosa *et al.* 1999; Cook *et al.* 2001). Thus, such integrated analyses may enable the construction of large phylogenies from morphology by use of datasets containing phylogenetic signal at multiple taxonomic levels.

**Figure 6:**
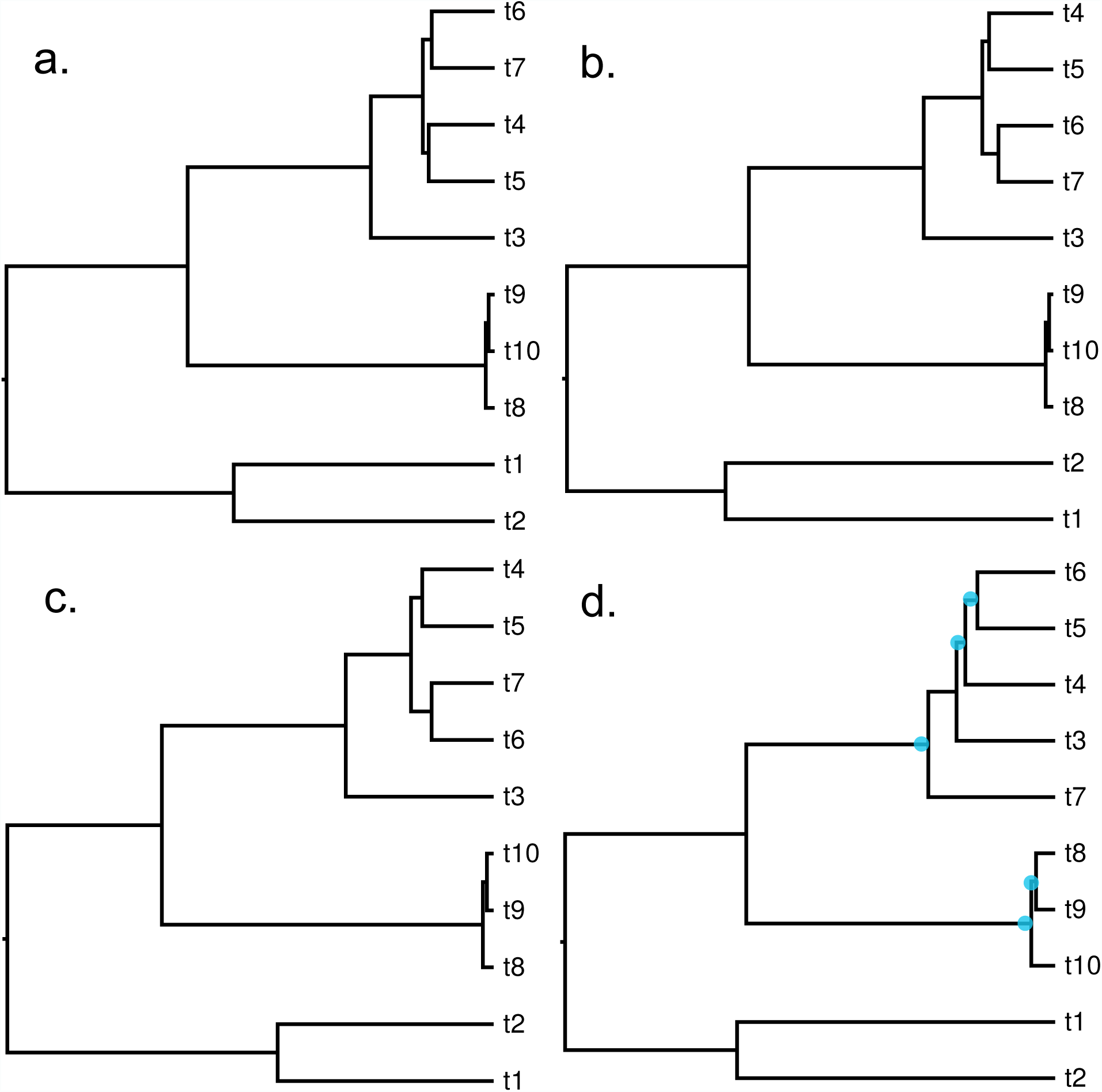
**a.** True tree. **b.** Tree estimated from 50 discrete and 50 continuous characters **c.** Tree estimated from 100 continuous characters simulated at rate 1.0 **d.** Tree estimated from 100 discrete characters simulated at rate 1.0. Blue dots signify incorrectly estimated bipartitions. The tree in panel b. was generated by randomly subsampling the matrices used to generate trees c. and d., and combining into a single matrix. This matrix was analysed in RevBayes. An example script is provided in the supplement.

Depending on the extent to which individual morphometric datasets are bounded and constrained in their evolution, analysis of continuous characters may help to increase phylogenetic information. Collecting morphometric measurements in many dimensions may enable the assembly of datasets that are large in size compared to those comprised of discrete characters alone. Although large collections of morphometric measurements may be strongly covarying, analysis of the correlation structure of such datasets, as mentioned above, would enable correction for biased signal and may reveal additional phylogenetic information. This would signify a more data-scientific approach to morphological phylogenetics by enabling researchers to dissect signal present in large morphometric datasets rather than reconstruct relationships using carefully curated data matrices. Such a paradigm shift would bring morphological phylogenetics closer in spirit to phylogenomic studies and enable deeper biological inferences through co-estimation of species relationships and dynamics in trait evolution. This would provide a firm phylogenetic backing to morphometric studies, and potentially reinvigorate the field in a similar way to the previous merging of phylogenetics and genomics. Improved ability to infer phylogeny among fossil taxa would also benefit molecular phylogenetics because the incorporation of fossils into total evidence matrices can improve both inference of molecular dates and alleviate long branch attraction (Huelsenbeck 1991; Wiens 2005; Ronquist *et al.* 2012). Though further study is needed to measure the expected phylogenetic information content of both continuous and discrete traits, all of the points discussed above should urge palaeontologists to give greater consideration to continuous traits in phylogenetic analysis of evolutionary patterns and relationships. This may improve efficiency in the use of hard-won palaeontological data by maximizing the amount of information gleaned from specimens and transform the field by facilitating new lines of questioning in palaeobiology.

And despite this optimistic tone, it should be noted that major work is still needed to provide deeper understanding of the behaviour of continuous trait models when used to infer phylogeny. It will be also important to gain a better understanding of expected empirical properties of continuous and discrete characters. As is shown here, discrete and continuous characters perform equally well when phylogenetic constraint is held constant, but there still lacks a clear characterisation of the relative expected constraint found in empirical datasets. As such, further work will be necessary to develop knowledge of the relative phylogenetic information content expressed across data types.

Moving forward, several extensions to existing Gaussian trait models should be explored. For example, further work is needed to determine the extent and distribution of rate heterogeneity between sites in continuous alignments. Since its presence has been well documented in molecular and discrete morphological data, it is likely that such rate heterogeneity is present in continuous measurements, and should be accommodated in empirical studies. Since traits can evolve under a broad range of processes, the fit of alternative models of continuous character evolution to empirical data and their adequacy in describing variation among them should also be examined.

### Is Mk a reasonable model for discrete character evolution?

Although likelihood approaches making use of the Mk model have been increasingly adopted in morphological phylogenetics, it is unclear whether it provides a reasonable approximation of the evolutionary process. Although there are explicit theoretical links between Markov substitution models and population genetic processes (Jukes and Cantor 1969), such theory does not exist in morphology. It should also be noted that molecular data are rarely modelled using the single parameter Jukes-Cantor model, with more complex generalisations typically preferred (Felsenstein 1981; Tavaré 1986). More sophisticated Markov processes can in principle be applied to morphological data, though this is rarely done. Nonetheless, MrBayes and RAxML implement HKY and General Time Reversible models, respectively, that can be applied to data with varying numbers of states (Ronquist and Huelsenbeck 2003; Stamatakis 2006). More work is needed to examine the adequacy of the Mk model in describing discrete character evolution. Such work will guide dataset assembly and the development of new model extensions. This is especially important in total-evidence tip dating methods employing Mk, as poor branch length estimates may weaken the ability to infer branching times. Although presenting a unique set of challenges, the use of continuous characters may alleviate some of issues concerning model misspecification. Models describing their change have been demonstrated to provide a reasonable description of character change resulting from several different microevolutionary processes (Hansen and Martins 1996). Further work is needed to address the relative adequacy of discrete and continuous trait models in describing the evolution of phenotypic data. In light of the results presented here, I suggest that continuous trait models be favoured in phylogenetic analysis in cases where morphological variation can be described quantitatively. Moving forward, deeper insight concerning the behaviour and adequacy of both discrete and continuous character models will enable increasingly powerful inferences to be drawn from morphological data. These issues will be of critical importance as advances in data collection and fossil evidence usher in an age of unprecedented discovery in morphological phylogenetics.

## Acknowledgements

I would like to thank JW Brown, SA Smith, GW Stull for helpful discussion of the manuscript. I would also like to thank DL Rabosky, AA King, EL Ionides, and M Friedman for sharing their thoughts on the subject and providing valuable feedback. Finally, I thank LJ Harmon, Thomas Guillerme, and two anonymous reviewers for their thoughtful comments which greatly benefited the manuscript.

---

**Figure S1:**
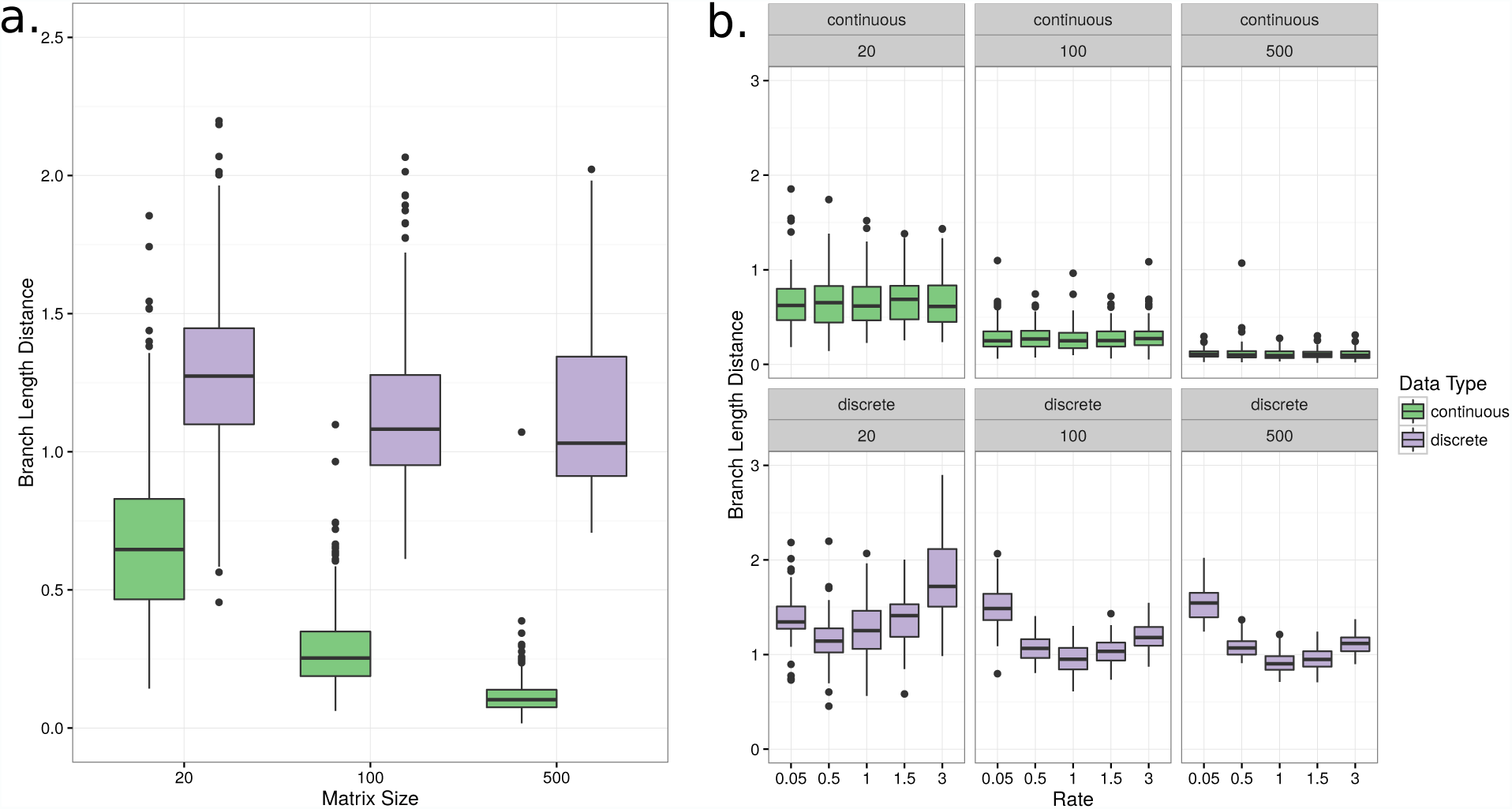
Branch length distance (BLD) across trees estimated from independently evolving continuous characters. **a.** BLD averaged across all rates except for the highest rate category, which resulted in the highest error when inferring under Mk. **b.** BLD across all matrix sizes and rates.

## References

Adams, D. C. and Felice, R. N. 2014. Assessing trait covariation and morphological integration on phylogenies using evolutionary covariance matrices. PloS One, 9(4): e94335.

Andersson, L., Haley, C. S., Ellegren, H., Knott, S. A., Johansson, M., Andersson, K.,Andersson-Eklund, L., Edfors-Lilja, I., Fredholm, M., Hansson, I., et al. 1994. Genetic mapping of quantitative trait loci for growth and fatness in pigs. Science, 263(5154): 1771–1775.

Arcila, D., Pyron, R. A., Tyler, J. C., Ortí, G., and Betancur-R, R. 2015. An evaluation of fossil tip-dating versus node-age calibrations in tetraodontiform fishes (teleostei: Percomorphaceae). Mol. Phylogenet. Evol., 82: 131–145.

Beaulieu, J. M., Jhwueng, D.-C., Boettiger, C., and O’Meara, B. C. 2012. Modeling stabilizing selection: expanding the ornstein–uhlenbeck model of adaptive evolution. Evolution, 66(8): 2369–2383.

Bloch, J. I. and Boyer, D. M. 2002. Grasping primate origins. Science, 298(5598): 1606–1610.

Bouckaert, R., Heled, J., Kühnert, D., Vaughan, T., Wu, C.-H., Xie, D., Suchard, M. A., Rambaut, A., and Drummond, A. J. 2014. Beast 2: a software platform for bayesian evolutionary analysis. PLoS Comput. Biol., 10(4): e1003537.

Brazeau, M. D. 2011. Problematic character coding methods in morphology and their effects. Biological Journal of the Linnean Society, 104(3): 489–498.

Burke, A. C., Nelson, C. E., Morgan, B. A., and Tabin, C. 1995. Hox genes and the evolution of vertebrate axial morphology. Development, 121(2): 333–346.

Butler, M. A. and King, A. A. 2004. Phylogenetic comparative analysis: a modeling approach for adaptive evolution. Am. Nat., 164(6): 683–695.

Cavalli-Sforza, L. L. and Edwards, A. W. 1967. Phylogenetic analysis. models and estimation procedures. Am. J. Hum. Genet., 19(3): 233.

Chang, J. and Alfaro, M. 2015a. Crowdsourced morphometric data are as accurate as traditionally collected data in 7 ray-finned fish families. Integr. Comp. Biol., 55: E28–E28.

Chang, J. and Alfaro, M. E. 2015b. Crowdsourced geometric morphometrics enable rapid large-scale collection and analysis of phenotypic data. Methods Ecol. Evol.

Chappill, J. A. 1989. Quantitative characters in phylogenetic analysis. Cladistics, 5(3): 217–234.

Cohn, M. J. and Tickle, C. 1999. Developmental basis of limblessness and axial patterning in snakes. Nature, 399(6735): 474.

Cook, C. E., Smith, M. L., Telford, M. J., Bastianello, A., and Akam, M. 2001. Hox genes and the phylogeny of the arthropods. Curr. Biol., 11(10): 759–763.

Dávalos, L. M., Velazco, P. M., Warsi, O. M., Smits, P. D., and Simmons, N. B. 2014. Integrating incomplete fossils by isolating conflicting signal in saturated and non-independent morphological characters. Syst. Biol., 63(4): 582–600.

De Rosa, R., Grenier, J. K., Andreeva, T., Cook, C. E., Adoutte, A., Akam, M., Carroll, S. B., and Balavoine, G. 1999. Hox genes in brachiopods and priapulids and protostome evolution. Nature, 399(6738): 772–776.

Dembo, M., Matzke, N. J., Mooers, A. Ø., and Collard, M. 2015. Bayesian analysis of a morphological supermatrix sheds light on controversial fossil hominin relationships. Proc. R. Soc. B, 282(1812): 20150943.

Donoghue, M. J. and Ree, R. H. 2000. Homoplasy and developmental constraint: A model and an example from plants 1. Amer. Zool., 40(5): 759–769.

Farris, J. S. 1990. Phenetics in camouflage. Cladistics, 6(1): 91–100.

Felsenstein, J. 1973. Maximum-likelihood estimation of evolutionary trees from continuous characters. Am. J. Hum. Genet., 25(5): 471.

Felsenstein, J. 1981. Evolutionary trees from dna sequences: a maximum likelihood approach. J. Mol. Evol., 17(6): 368–376.

Felsenstein, J. 1985. Phylogenies and the comparative method. Am. Nat., 125(1): 1–15.

Frary, A., Nesbitt, T. C., Frary, A., Grandillo, S., Van Der Knaap, E., Cong, B., Liu, J., Meller, J., Elber, R., Alpert, K. B., et al. 2000. fw2. 2: a quantitative trait locus key to the evolution of tomato fruit size. Science, 289(5476): 85–88.

Gingerich, P. D. 1993. Quantification and comparison of evolutionary rates. Am. J. Sci., 293(A): 453–478.

Goloboff, P. A., Mattoni, C. I., and Quinteros, A. S. 2006. Continuous characters analyzed as such. Cladistics, 22(6): 589–601.

Guillerme, T. and Cooper, N. 2016. Effects of missing data on topological inference using a total evidence approach. Molecular phylogenetics and evolution, 94: 146–158.

Hansen, T. F. 1997. Stabilizing selection and the comparative analysis of adaptation. Evolution, pages 1341–1351.

Hansen, T. F. and Martins, E. P. 1996. Translating between microevolutionary process and macroevolutionary patterns: the correlation structure of interspecific data. Evolution, pages 1404–1417.

Hauser, D. L. and Presch, W. 1991. The effect of ordered characters on phylogenetic reconstruction. Cladistics, 7(3): 243–265.

Hawkins, J. A., Hughes, C. E., and Scotland, R. W. 1997. Primary homology assessment, characters and character states. Cladistics, 13(3): 275–283.

Hill, W. and Robertson, A. 1968. Linkage disequilibrium in finite populations. Theor. Appl. Genet., 38(6): 226–231.

Hillis, D. and Huelsenbeck, J. 1992. Signal, noise, and reliability in molecular phylogenetic analyses. J. Hered., 83(3): 189–195.

Höhna, S., Landis, M. J., Heath, T. A., Boussau, B., Lartillot, N., Moore, B. R., Huelsenbeck, J. P., and Ronquist, F. 2016. Revbayes: Bayesian phylogenetic inference using graphical models and an interactive model-specification language. Syst. Biol., 65(4): 726–736.

Huelsenbeck, J. P. 1991. When are fossils better than extant taxa in phylogenetic analysis? Syst. Biol., 40(4): 458–469.

Hunt, G. J., Guzmán-Novoa, E., Fondrk, M. K., and Page, R. E. 1998. Quantitative trait loci for honey bee stinging behavior and body size. Genetics, 148(3): 1203–1213.

Jukes, T. H. and Cantor, C. R. 1969. Evolution of protein molecules. Mammalian Protein Metab., 3(21): 132.

Kirk, E. C., Cartmill, M., Kay, R. F., and Lemelin, P. 2003. Comment on” grasping primate origins”. Science, 300(5620): 741–741.

Klopfstein, S., Vilhelmsen, L., and Ronquist, F. 2015. A nonstationary markov model detects directional evolution in hymenopteran morphology. Systematic biology, 64(6): 1089–1103.

Kuhner, M. K. and Felsenstein, J. 1994. A simulation comparison of phylogeny algorithms under equal and unequal evolutionary rates. Mol. Biol. Evol., 11(3): 459–468.

Landis, M. J., Schraiber, J. G., and Liang, M. 2013. Phylogenetic analysis using lévy processes: finding jumps in the evolution of continuous traits. Syst. Biol., 62(2): 193–204.

Lee, M. S. and Palci, A. 2015. Morphological phylogenetics in the genomic age. Curr. Biol., 25(19): R922–R929.

Lee, M. S. and Worthy, T. H. 2012. Likelihood reinstates archaeopteryx as a primitive bird. Biol. Lett., 8(2): 299–303.

Lee, M. S., Soubrier, J., and Edgecombe, G. D. 2013. Rates of phenotypic and genomic evolution during the cambrian explosion. Curr. Biol., 23(19): 1889–1895.

Lee, M. S., Cau, A., Naish, D., and Dyke, G. J. 2014. Morphological clocks in paleontology, and a mid-cretaceous origin of crown aves. Syst. Biol., 63(3): 442–449.

Lewis, P. O. 2001. A likelihood approach to estimating phylogeny from discrete morphological character data. Syst. Biol., 50(6): 913–925.

Livezey, B. C. and Zusi, R. L. 2007. Higher-order phylogeny of modern birds (theropoda, aves: Neornithes) based on comparative anatomy. ii. analysis and discussion. Zool. J. Linnean Soc., 149(1): 1–95.

Lynch, M., Walsh, B., et al. 1998. Genetics and analysis of quantitative traits, volume 1. Sinauer Sunderland, MA.

McVean, G. 2007. The structure of linkage disequilibrium around a selective sweep. Genetics, 175(3): 1395–1406.

Nylander, J. A., Ronquist, F., Huelsenbeck, J. P., and Nieves-Aldrey, J. 2004. Bayesian phylogenetic analysis of combined data. Syst. Biol., 53(1): 47–67.

O’leary, M. A., Bloch, J. I., Flynn, J. J., Gaudin, T. J., Giallombardo, A., Giannini, N. P., Goldberg, S. L., Kraatz, B. P., Luo, Z.-X., Meng, J., et al. 2013. The placental mammal ancestor and the post–k-pg radiation of placentals. Science, 339(6120): 662–667.

O’Reilly, J. E., Puttick, M. N., Parry, L., Tanner, A. R., Tarver, J. E., Fleming, J., Pisani, D., and Donoghue, P. C. 2016. Bayesian methods outperform parsimony but at the expense of precision in the estimation of phylogeny from discrete morphological data. Biol. Lett., 12(4): 20160081.

Pagel, M. 1994. Detecting correlated evolution on phylogenies: a general method for the comparative analysis of discrete characters. Proc. R. Soc. B, 255(1342): 37–45.

Palaisa, K., Morgante, M., Tingey, S., and Rafalski, A. 2004. Long-range patterns of diversity and linkage disequilibrium surrounding the maize y1 gene are indicative of an asymmetric selective sweep. Proc. Natl. Acad. Sci. U. S. A., 101(26): 9885–9890.

Pattinson, D. J., Thompson, R. S., Piotrowski, A. K., and Asher, R. J. 2014. Phylogeny, paleontology, and primates: do incomplete fossils bias the tree of life? Syst. Biol., 64: 169–186.

Pennell, M. W. and Harmon, L. J. 2013. An integrative view of phylogenetic comparative methods: connections to population genetics, community ecology, and paleobiology. Ann. N. Y. Acad. Sci., 1289(1): 90–105.

Pimentcl, R. A. and Riggins, R. 1987. The nature of cladistic data. Cladistics, 3(3): 201–209.

Pleijel, F. 1995. On character coding for phylogeny reconstruction. Cladistics, 11(3): 309–315.

Puttick, M. N., O’Reilly, J. E., Tanner, A. R., Fleming, J. F., Clark, J., Holloway, L., Lozano-Fernandez, J., Parry, L. A., Tarver, J. E., Pisani, D., and Donoghue, P. C. J. 2017. Uncertain-tree: discriminating among competing approaches to the phylogenetic analysis of phenotype data. Proc. R. Soc. B, 284(1846).

R Core Team 2016. R: A Language and Environment for Statistical Computing. R Foundation for Statistical Computing, Vienna, Austria.

Rae, T. C. 1998. The logical basis for the use of continuous characters in phylogenetic systematics. Cladistics, 14(3): 221–228.

Rambaut, A. and Drummond, A. 2013. Treeannotator v1. 7.0. Available as part of the BEAST package at http://beast.bio.ed.ac.uk.

Reich, D. E., Cargill, M., Bolk, S., Ireland, J., Sabeti, P. C., Richter, D. J., Lavery, T., Kouyoumjian, R., Farhadian, S. F., Ward, R., et al. 2001. Linkage disequilibrium in the human genome. Nature, 411(6834): 199–204.

Revell, L. J. 2012. phytools: an r package for phylogenetic comparative biology (and other things). Methods Ecol. Evol., 3(2): 217–223.

Revell, L. J., Harmon, L. J., and Collar, D. C. 2008. Phylogenetic signal, evolutionary process,and rate. Syst. Biol., 57(4): 591–601.

Robinson, D. F. and Foulds, L. R. 1981. Comparison of phylogenetic trees. Math. Biosci., 53(1-2): 131–147.

Ronquist, F. and Huelsenbeck, J. P. 2003. Mrbayes 3: Bayesian phylogenetic inference under mixed models. Bioinformatics, 19(12): 1572–1574.

Ronquist, F., Klopfstein, S., Vilhelmsen, L., Schulmeister, S., Murray, D. L., and Rasnitsyn, A. P. 2012. A total-evidence approach to dating with fossils, applied to the early radiation of the hymenoptera. Syst. Biol., 61(6): 973–999.

Ronquist, F., Lartillot, N., and Phillips, M. J. 2016. Closing the gap between rocks and clocks using total-evidence dating. Phil. Trans. R. Soc. B, 371(1699): 20150136.

Schlenke, T. A. and Begun, D. J. 2004. Strong selective sweep associated with a transposon insertion in drosophila simulans. Proc. Natl. Acad. Sci. U. S. A., 101(6): 1626–1631.

Scotland, R. and Pennington, R. T. 2000. Homology and systematics: coding characters for phylogenetic analysis. CRC Press.

Scotland, R. W., Olmstead, R. G., and Bennett, J. R. 2003. Phylogeny reconstruction: the role of morphology. Syst. Biol., 52(4): 539–548.

Simões, T. R., Caldwell, M. W., Palci, A., and Nydam, R. L. 2017. Giant taxon-character matrices: quality of character constructions remains critical regardless of size. Cladistics, 33(2): 198–219.

Smith, U. E. and Hendricks, J. R. 2013. Geometric morphometric character suites as phylogenetic data: extracting phylogenetic signal from gastropod shells. Syst. Biol., 62: 366–385.

Stamatakis, A. 2006. Raxml-vi-hpc: maximum likelihood-based phylogenetic analyses with thousands of taxa and mixed models. Bioinformatics, 22(21): 2688–2690.

Sukumaran, J. and Holder, M. T. 2010. Dendropy: a python library for phylogenetic computing. Bioinformatics, 26(12): 1569–1571.

Tavaré, S. 1986. Some probabilistic and statistical problems in the analysis of dna sequences. Lectures on mathematics in the life sciences, 17: 57–86.

Thiele, K. 1993. The holy grail of the perfect character: the cladistic treatment of morphometric data. Cladistics, 9(3): 275–304.

Upchurch, P. 1995. The evolutionary history of sauropod dinosaurs. Philos. Trans. R. Soc. B, 349(1330): 365–390.

Uyeda, J. C., Caetano, D. S., and Pennell, M. W. 2015. Comparative analysis of principal components can be misleading. Syst. Biol., 64(4): 677–689.

Valdar, W., Solberg, L. C., Gauguier, D., Burnett, S., Klenerman, P., Cookson, W. O., Taylor, M. S., Rawlins, J. N. P., Mott, R., and Flint, J. 2006. Genome-wide genetic association of complex traits in heterogeneous stock mice. Nat. Genet., 38(8): 879–887.

Wagner, P. J. 2012. Modelling rate distributions using character compatibility: implications for morphological evolution among fossil invertebrates. Biol. Lett., 8(1): 143–146.

Wickham, H. 2016. ggplot2: elegant graphics for data analysis. Springer.

Wiens, J. J. 2001. Character analysis in morphological phylogenetics: problems and solutions. Syst. Biol., 50(5): 689–699.

Wiens, J. J. 2005. Can incomplete taxa rescue phylogenetic analyses from long-branch attraction? Syst. Biol., 54(5): 731–742.

Wiens, J. J., Kuczynski, C. A., Townsend, T., Reeder, T. W., Mulcahy, D. G., and Sites, J. W. 2010. Combining phylogenomics and fossils in higher-level squamate reptile phylogeny: molecular data change the placement of fossil taxa. Syst. Biol., 59(6): 674–688.

Wilkinson, D. G., Bhatt, S., Cook, M., Boncinelli, E., and Krumlauf, R. 1989. Segmental expression of hox-2 homoeobox-containing genes in the developing mouse hindbrain. Nature, 341: 405–409.

Wilkinson, M. 1995. A comparison of two methods of character construction. Cladistics, 11(3): 297–308.

Wilson, J. A. and Sereno, P. C. 1998. Early evolution and higher-level phylogeny of sauropod dinosaurs. J. Vert. Paleontol., 18(S2): 1–79.

Wood, H. M., Matzke, N. J., Gillespie, R. G., and Griswold, C. E. 2012. Treating fossils as terminal taxa in divergence time estimation reveals ancient vicariance patterns in the palpimanoid spiders. Syst. Biol., 62(2): 264–282.

Wright, A. M. and Hillis, D. M. 2014. Bayesian analysis using a simple likelihood model outperforms parsimony for estimation of phylogeny from discrete morphological data. PLoS One, 9(10): e109210.

Wright, A. M., Lloyd, G. T., and Hillis, D. M. 2015. Modeling character change heterogeneity in phylogenetic analyses of morphology through the use of priors. Syst. Biol., page syv122.

Yang, Z. 1998. On the best evolutionary rate for phylogenetic analysis. Syst. Biol., 47(1): 125–133.

